# Confocal imaging capacity on a widefield microscope using a spatial light modulator

**DOI:** 10.1101/2020.12.03.409581

**Authors:** Yao L. Wang, Noa W. F. Grooms, Sabrina C. Civale, Samuel H. Chung

## Abstract

Confocal microscopes can reject out-of-focus and scattered light; however, widefield microscopes are far more common in biological laboratories due to their accessibility and lower cost. We report confocal imaging capacity on a widefield microscope by adding a spatial light modulator (SLM) and utilizing custom illumination and acquisition methods. We discuss our illumination strategy and compare several procedures for postprocessing the acquired image data. We assessed the performance of this system for rejecting out-of-focus light by comparing images taken using our widefield microscope, our SLM-enhanced setup, and a commercial confocal microscope. The optical sectioning capability, assessed on thin fluorescent film, was 0.85 ± 0.04 μm for our SLM-enhanced setup and 0.68 ± 0.04 μm for a confocal microscope, while a widefield microscope exhibited no sectioning capability. We demonstrate our setup by imaging the same set of neurons in *C. elegans* on widefield, SLM, and confocal microscopes. SLM enhancement greatly reduces background from the cell body, allowing visualization of dim fibers nearby. Our SLM-enhanced setup identified 93% of the dim neuronal fibers seen in confocal images while a widefield microscope only identified 48% of the same fibers. Our microscope add-on represents a very simple (2-component) and inexpensive (<$600) approach to enable widefield microscopes to optically section thick samples.

## Introduction

Widefield epifluorescence microscopes have driven numerous advances in the biological sciences and are ubiquitous in laboratories. Despite their powerful capabilities, broad accessibility, and relatively low cost, widefield microscopes cannot exclude out-of-focus or scattered light. In sparsely-populated or sparsely-labelled samples, this weakness has relatively minor impact: The illumination light is focused onto the plane being observed, so out-of-focus objects are illuminated by a lower intensity of light and in-focus objects are more likely to dominate images. The light from these out-of-focus objects, however, is not excluded. It remains diffusely in the image and interferes with imaging. This weakness has spurred the development of scanning techniques such as confocal microscopy, which can reject both out-of-focus and scattered light [1]. The key component in confocal microscopes is a pinhole in the emission path, which excludes out-of-focus light. Point scanning, in combination with the pinhole, also excludes scattered light as only imaging light from the excited location is included in the final image. Exclusion of unwanted light allows confocal to have significantly better resolution and optical sectioning ability than widefield microscopy. The disadvantages of confocal microscopy are greatly increased complexity, moving parts, requirement of synchronization, and consequently, increased cost.

In our study, we developed simple and inexpensive methods to reduce and exclude scattered and out-of-focus light in a standard widefield epifluorescence microscope. In our approach, we spatially modulate the illumination light and postprocess captured images. Our technique capitalizes on pixelated arrays, such as the spatial light modulator (SLM) and the digital micromirror device (DMD), which modulate the intensity of transmitted and/or reflected light. A transmissive SLM between two crossed polarizers selectively transmits varying intensities of light through each array element. Many projectors use these SLM devices to display an image using incoherent light from a projector bulb. As previously reviewed [2], SLMs and DMDs at the field or aperture stop can control the spatial distribution of the excitation light [3], select the sample location that is observed [4], exclude out-of-focus light [5, 6], and perform structured illumination [7]. Pixelated arrays represent a flexible and cost-effective approach [8] to spatially modulate a light distribution for multiple applications.

Extending the efforts of prior studies, here we show that a transmissive SLM can selectively illuminate locations in the sample with < 500 nm resolution when placed at the field stop of an epifluorescence microscope. Together with minimal postprocessing, we demonstrate optical sectioning (*i.e.*, confocal) capability on a typical widefield microscope using an inexpensive SLM add-on. We characterize images taken on our setup and compare them to images from widefield and confocal microscopes. We demonstrate significantly clearer *in vivo* imaging of neurons in *C. elegans* compared to typical widefield microscopy.

## Results

### Basic configuration and concept

The modern widefield epifluorescence microscope (see Fig. 1a) is centered around an objective that focuses Köhler illumination light onto a sample and captures emission light from the sample. The emission light is imaged by a tube lens, often onto a camera array. Many widefield microscopes have additional optics in the excitation beampath, represented by two lenses in Fig. 1a, that shape and optimize the excitation light. These optics create two planes, the field and aperture (not shown) stops, where masks can be inserted to control the extent and angle of the excitation, respectively.

**Figure 1.**
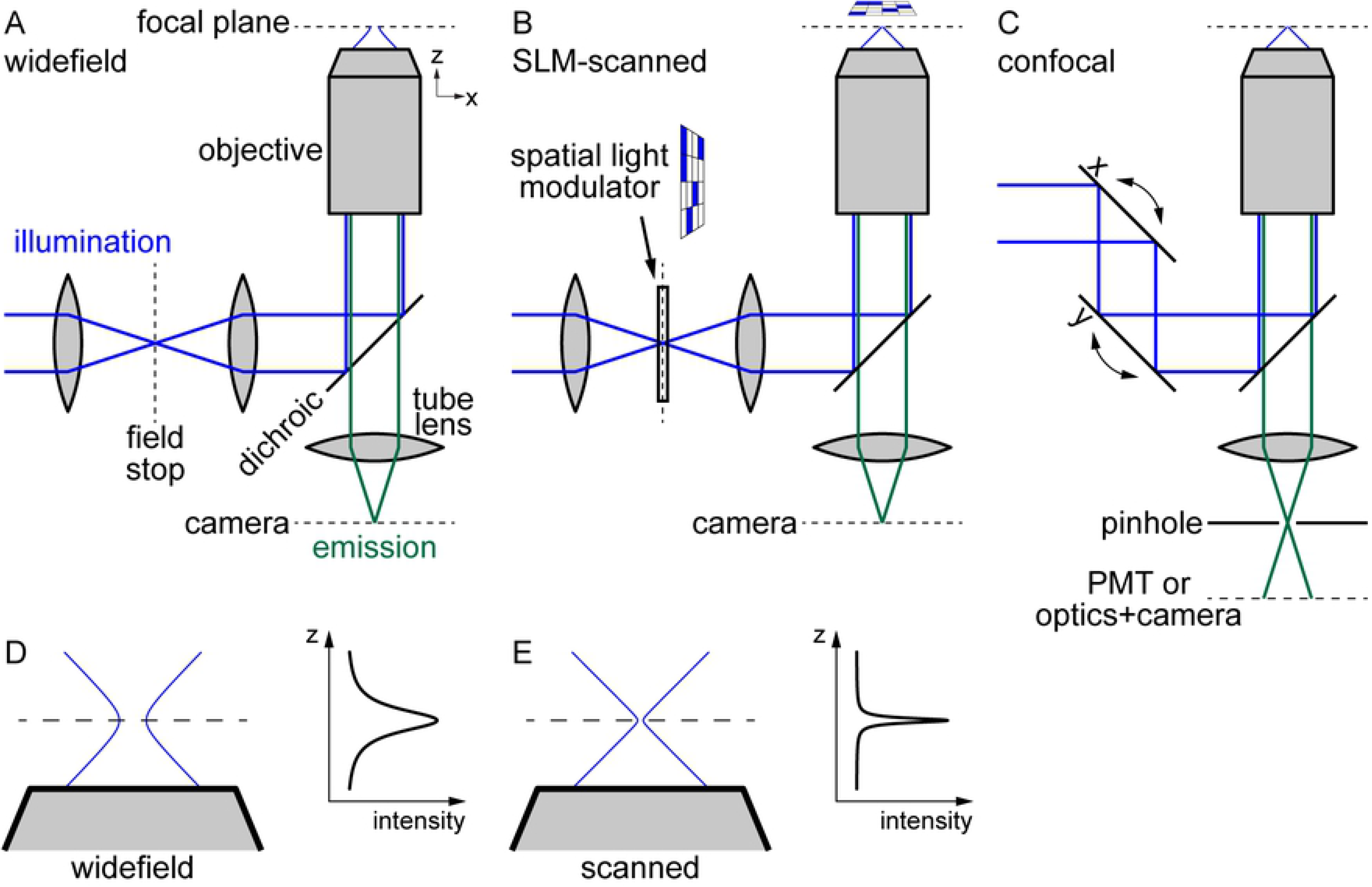
Optics and light beampaths for microscope setups. (a) Widefield illumination optics create field stop plane, whose light distribution is projected to focal plane in sample via objective. Camera images emission light from focal plane. (b) Introduction of SLM at field stop allows arbitrary illumination patterns at focal plane. (c) Confocal microscope optics scan illumination beam and utilize pinhole to exclude out-of-focus emission light. (d) Widefield axial illumination profile. Wide intensity peak around focal plane. (e) Scanned axial illumination profile. Narrow intensity peak around focal plane.

In contrast to the simplicity of widefield microscopes, confocal microscopes (simplified diagram in Fig. 1c) utilize complex sets of optics in both the excitation and emission beampaths. The excitation optics sweep the excitation light across the sample to image each location in turn. The optics depend on the type of confocal microscope: Laser excitation is typically scanned with mirrors [9]. Spinning disk excitation illuminates multiple, distant points in the sample simultaneously by passing illumination light through a Nipkow disk containing multiple holes [10]. In general, the emission optics include the key optic that distinguishes confocal from widefield microscopy: a pinhole positioned at a conjugate focal plane in the emission beampath. The pinhole strongly filters out light that does not originate from the focal plane, allowing clear imaging of a two-dimensional slice in the bulk without stray light from other depths (*e.g.*, optical sectioning). Because the emission light comes from different locations in the sample, it can exit the objective at various angles and must be “descanned”. Practically, emission light is typically counterpropagated through the same optics used to scan the excitation beam (not shown in figure for simplicity). It is then spectrally separated by a dichroic mirror and filtered by the pinhole prior to detection by a photomultiplier tube (PMT) for laser scanning confocal or by a camera array for spinning disk confocal.

In our setup, the SLM is inserted into the field stop position, so that the transmitted light distribution is projected at the focal plane in the sample (see Fig. 1b). The key functional difference between widefield and SLM scanned illumination is that SLM scanning concentrates the illumination to the focal plane. The underlying mechanism is shown in Fig. 1de. The different beam profiles of the widefield and SLM scanned configurations leads to different intensity profiles in z. The intensity is I = P / A = P / πr^2^, where I, P, A, and r are the optical intensity, optical power, beam cross-sectional area, and beam radius, respectively. The widefield illumination beam is purposely diffuse so that it illuminates the entire field of view at the focal plane. Because the focal area remains large at the focal plane (dotted line), the radius changes relatively little near the focal plane, and the intensity profile in the axial direction is relatively wide. Conversely, the SLM-scanned beam can illuminate a small region of the focal plane (corresponding to transmission through a single SLM element). Near the focal plane, the radius comes to a sharp minimum. Thus, assuming the same numerical aperture as the widefield configuration, the intensity profile in the axial direction is relatively narrow. By concentrating the illumination at the focal plane, we reduce the emission light from out-of-focus objects and enhance the ability of the microscope to optically section the sample, even without true confocal operation. As described below, we further achieve true confocal operation via postprocessing.

### Implementation: illumination and postprocessing

We utilize a transmissive SLM taken from a commercial projector, attached to a 3D printed mount and a 6-axis stage for precise alignment in the field stop (see Materials and Methods, Fig. 2a). One advantage of using a transmissive SLM is its high accessibility: no other modification to a widefield microscope is required for integration. However, the transmissive SLM has several weaknesses in such implementation. First, its transmission is low, making it challenging to obtain sufficient illumination power, particularly for dim samples, where it could potentially improve imaging the most. Second, the fill factor, or fractional area of the SLM that is actively modulated is low (see Fig. S1), typically slightly above 60%. This low fill factor is due to conduits at the periphery of each element to conduct signals that control the transmission of the active regions. The non-active region casts a shadow at the sample, making the final image pixelated. Even with these weaknesses, we still demonstrate greatly improved imaging by adding a transmissive SLM to our widefield microscope.

**Figure 2.**
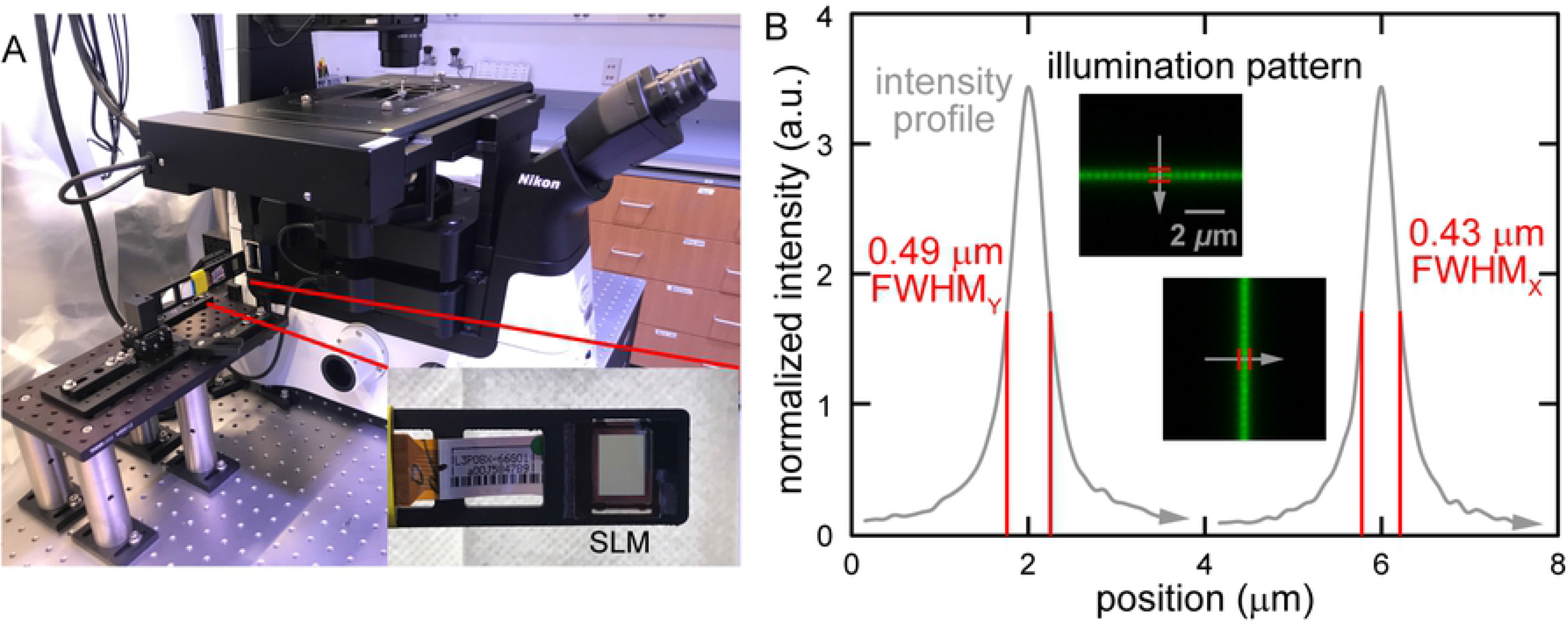
Illumination setup and resolution. (a) SLM on 3D-printed mount on 6-axis fine-positioning stage. SLM inserted into field stop of microscope. (b) Illumination resolution determined using fluorescent dye thin film.

We characterized the resolution of the SLM illumination by transmitting only through single lines of SLM elements and illuminating a thin film of fluorescent dye. We measured the illumination resolution as 0.43 μm in the horizontal and 0.49 μm in the vertical directions (see Fig. 2b), or an aspect ratio of 0.88. The non-uniform resolution is due to a rectangular active region of our SLM elements. As shown in Fig. S1, we used a brightfield dissecting microscope to measure the aspect ratio of the active region as 109 to 127 pixels, or 0.86, producing the difference in x and y resolution observed. We expect a uniform aspect ratio and finer resolution with an improved SLM or a DMD.

Stray light arises from fluorescence that does not originate from the sample position under observation, whether inside or outside of the focal plane. Illuminating small areas sequentially and separating these illuminated areas (*i.e.*, sparse illumination) reduces stray light as it must propagate further to interfere with useful fluorescence. We illuminate separated areas in the sample by transmitting a dot array of single SLM elements (see Fig. 3a) [6, 11]. For clarity in our description, we refer to individual components of the SLM as “elements” and individual components of the camera as “pixels”. Empirically, we determined that separating transmitting elements by five non-transmitting elements (6 × 6 unit cell) eliminates most light interference between illuminated points (see Fig. 3b) while maximizing speed. The size of the unit cell can be adjusted to balance imaging speed and quality. To illuminate all locations in the field of view (FOV), the dot array spans the entire FOV and is raster scanned through all 36 positions in the unit cell. We acquire a sub-image at each position and post-process these 36 sub-images for each final image.

**Figure 3.**
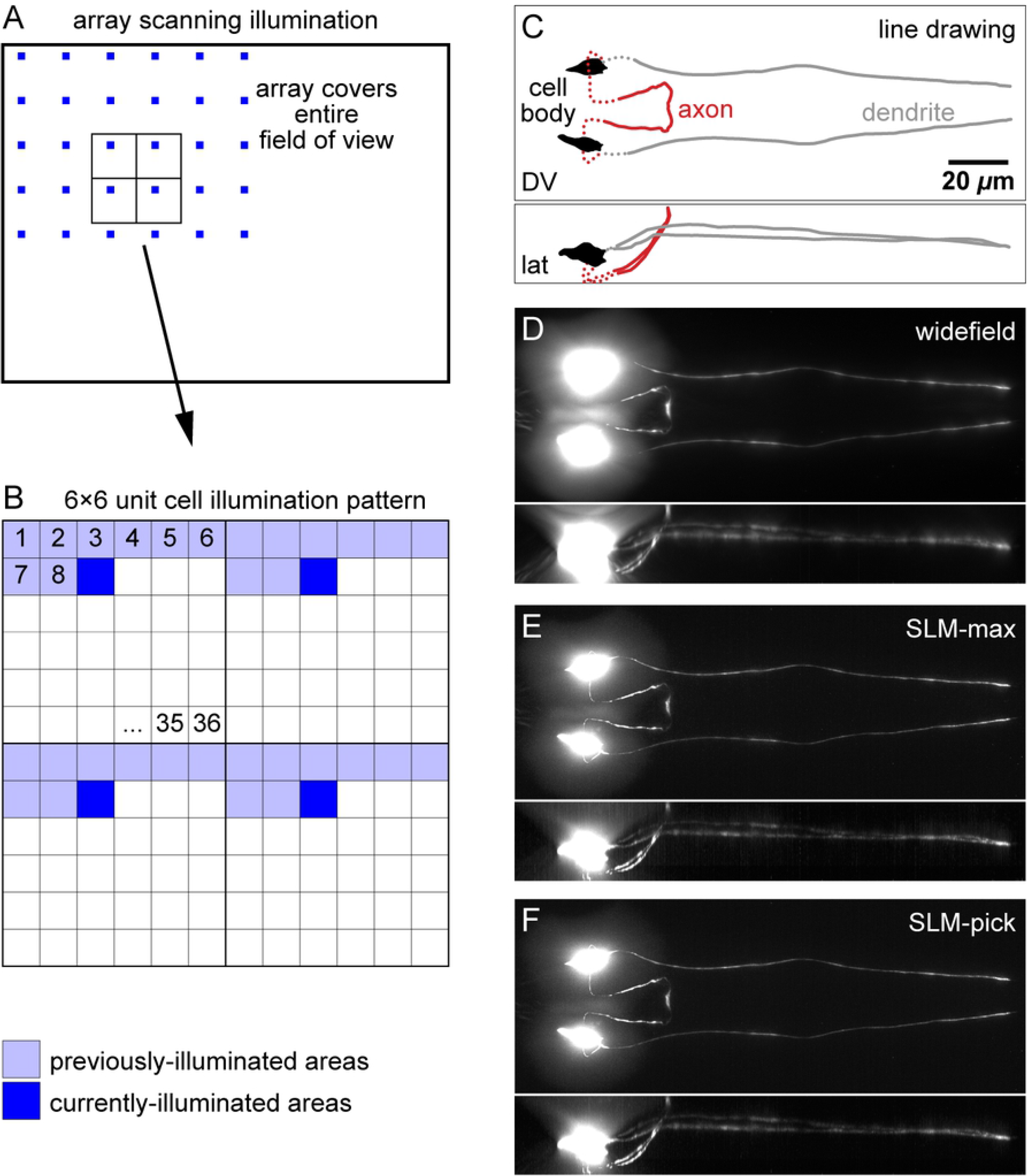
Illumination strategies and postprocessing. (a) Dot array transmitted by SLM illuminates sample. Array is raster scanned across FOV. (b) Expansion of inset in (a). Raster scanning moves single-element illumination through 36 locations of 6×6 unit cell. Sub-image of entire FOV is acquired for each of 36 illumination patterns. (c) Line drawing of ASJ neuron imaged. Dorsal-ventral view in xy direction (top), lateral view in xz direction (bottom). (d-f) Fluorescence images, obtained by maximum projections of 3D image. (d) Conventional widefield image. (e) SLM-max image. Pixel value in 3D image is maximum pixel value of 36 subimages. (f) SLM-pick image. Pixel value is in 3D image is pixel value when corresponding sample location is illuminated. Note significantly improved optical sectioning in the max and pick strategies, mostly visibly above and below the cell bodies.

Because we employed off-the-shelf SLMs, the area in the sample illuminated by a single SLM element is imaged by ~3.5 camera pixels. Along with the non-active region of transmission SLMs, this leads to some minor pixelation and aliasing in the final image (see insets in Fig. 2b). We expect to eliminate most of these artifacts by employing reflective DMDs, which have improved specifications. Performance will also greatly improve if the devices are designed for our microscope and camera.

We imaged the ASJ neuron in *C. elegans* (see Fig. 3c) to test three strategies for postprocessing images. We compared the resulting images to widefield images taken on the same microscope (see Fig. 3d, replicated in S2a). First, we summed the fluorescence values (“SLM-sum”) from the corresponding pixels in the 36 sub-images to calculate the intensity of each pixel in the final image (see Fig. S2b). Second, we took a maximum projection of the 36 sub-images, keeping the brightest of the 36 values of each pixel (“SLM-max”) as the intensity of the pixel in the final image (see Fig. 3e). In the third (“SLM-pick”), the intensity of the pixel in the final image is the pixel intensity in the sub-image during which the pixel was illuminated by the SLM (see Fig. 3f). As expected, the SLM-sum strategy produces an image that is very similar to a widefield image because we capture light while illuminating every position. The key difference is merely that light capture occurs sequentially in time rather than simultaneously as it does in widefield. The SLM-max strategy produced a final image that was slightly inferior to the pick strategy. However, the SLM-max strategy is simpler to implement than the pick strategy for crucial reasons. The SLM-pick strategy requires precise alignment of SLM elements to camera pixels in space and time: vertical location, horizontal location, and rotation around the optical axis as well as tight synchronization of the illumination and observation. Because SLM-sum and SLM-max strategies relax the alignment requirement, they are also significantly more robust to misalignment and instrument drift compared to the pick strategy. The SLM-max strategy relies on the brightest intensity at a location arising when that location is illuminated. Thus, the resulting image from SLM-max strategy approaches the SLM-pick strategy for sparsely-labelled samples but produces inferior results when bright structures are nearby, (*e.g.*, in an out-of-focus plane). In addition, the SLM-max strategy retains the brightest pixels and so generally leads to noisier images. The SLM-pick strategy captures fluorescent light from small, separated regions only while they are illuminated. This operation is similar to the operation of a pinhole in confocal microscopy. Thus, this strategy leads to images with the best rejection of out-of-focus and scattered light. The alignment requirement is readily achievable utilizing a high-precision stage (see Fig. 2a), and we use the SLM-pick strategy for the remainder of our study.

### Characterization on well-defined samples

We employed two methods for assessing the axial, or z-sectioning capability of our SLM-pick strategy compared to widefield and commercial confocal. First, we acquired 3D images of a thin fluorescent film. In agreement with prior results [5], axial profiles of widefield microscopy images do not exhibit a measurable peak, indicating minimal sectioning ability (see red curve in Fig. 4a). In contrast, the SLM-pick (purple curve) and confocal (blue curve) modalities show sharp peaks with full width at half maximum (FWHM) of 0.85 ± 0.04 μm and 0.68 ± 0.04 μm, respectively (see Fig. 4b). The experimental value of the SLM-pick FWHM matches well with a theoretical value of 0.83 μm, calculated from equation 4 of ref. [12]. This value is calculated with input parameters λ_em_ = 525 nm, n = 1.515, NA = 1.4, and PH = object-side pinhole diameter = image-side pinhole / magnification = 4 pixels * 6.5 μm/pixel / 60x = 0.43 μm. These data indicate the capability of SLM illumination and minimal postprocessing to produce images with optical sectioning. Second, we acquired 3D images of 6-μm fluorescent beads to mimic out-of-focus fluorescence from round cell bodies. Figure 4c shows transverse intensity profiles, and Fig. 4d shows axial intensity profiles through the center of an average image of 50 beads. To better highlight the impact bright cell bodies have on dim nearby fibers, we measured the full width at 10% max (FWTM). This value represents the approximate spatial range where objects with 10% the brightness of the bead (see dotted lines) are obscured by stray light from the bead. In the transverse direction (see Fig. 4e), the FWTM of the widefield, SLM, and confocal differ by < 15% due to the transverse confinement of focused light. In the axial direction (see Fig. 4f), the confocal FWTM is significantly smaller than the widefield FWTM because of optical sectioning. The SLM-generated imaging optical sectioning is about 30% greater than confocal but about 50% less than widefield.

**Figure 4.**
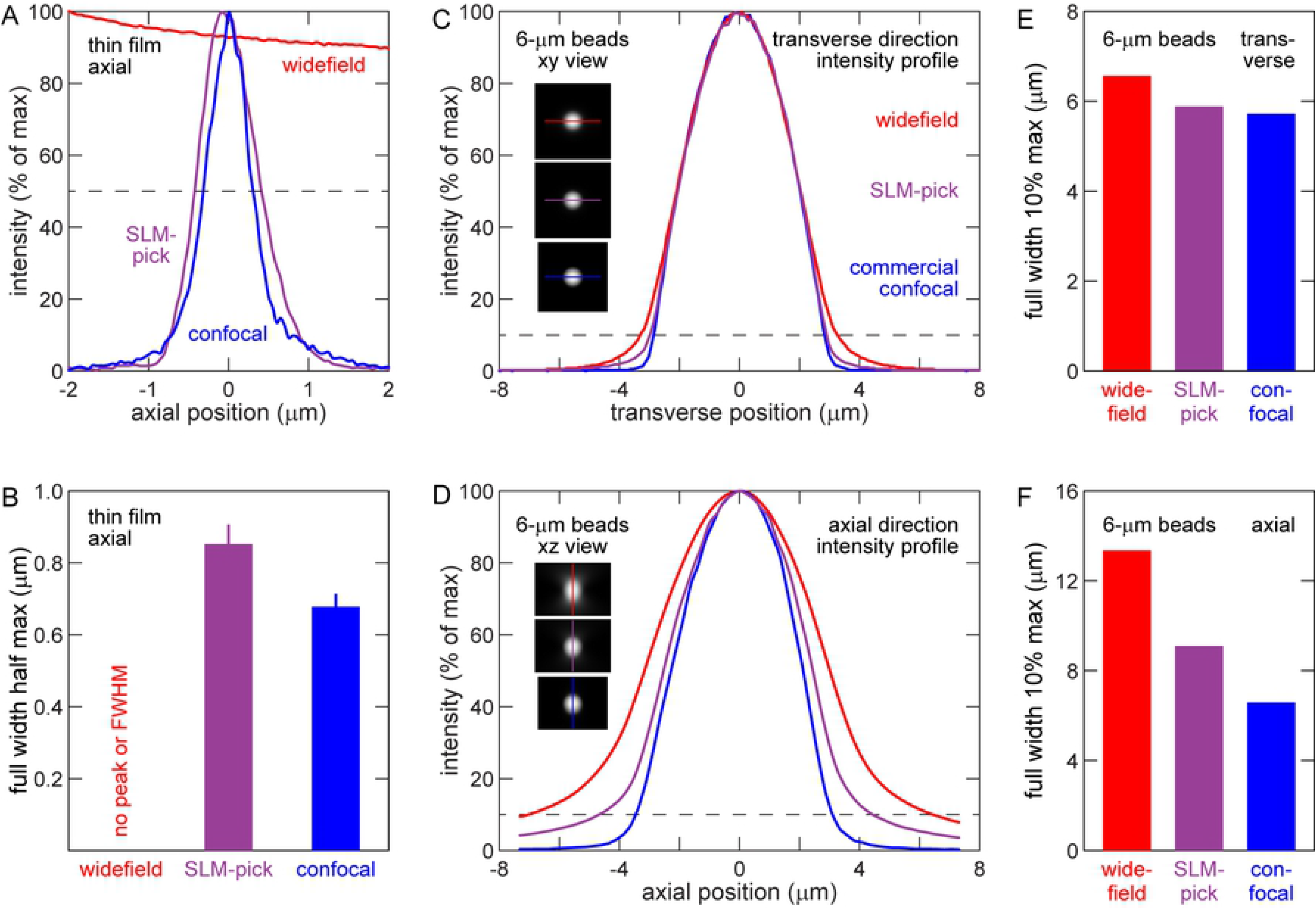
SLM improves sectioning capability compared to widefield imaging. (a) Intensity of light at axial positions around fluorescent thin film at *z* = 0. Dotted line at 50% maximum intensity. (b) FWHM of data in (a). *n* = 20 locations. Note optical sectioning capability conferred by SLM. (c) Average transverse intensity profile of 6-μm fluorescent beads. Dotted line at 10% maximum intensity. Maximum projection of average bead images shown in insets, with profile line. (d) Average axial intensity profile of 6-μm fluorescent beads. Maximum projection of average bead images shown in insets, with profile line. (e) Transverse full width at 10% maximum intensity in (c). (f) Axial full width at 10% maximum intensity in (d). Note improved confinement of fluorescence to true bead extent in SLM imaging compared to widefield, particularly in axial direction. *n* = 85 (widefield), 90 (SLM-pick), and 63 (confocal) beads.

While the SLM-pick strategy improves optical sectioning, we did not observe enhancement of resolution by either the SLM-max or SLM-pick strategies. As shown in Supp. Table S1, the transverse and axial FWHM of sub-diffraction-limit beads is unaffected by use of the SLM. This is because the resolution of our illumination (~0.45 μm) is significantly greater than the diffraction limit (~0.2 μm).

### Demonstration *in vivo*

Utilizing the SLM pick strategy, we imaged two types of samples to demonstrate our technique’s capabilities for *in vivo* imaging and compare performance with widefield and confocal imaging. First, to demonstrate enhanced imaging capabilities at high resolution, we imaged a *C. elegans* strain with a fluorescently-labelled class of neurons called the amphids, whose neuronal fibers are tightly bundled (see Fig. 5a). Widefield imaging has difficulty clearly resolving individual neuronal fibers due to stray light from nearby structures (see Fig. 5b and insets). As they reject stray light, SLM-pick (Fig. 5c) and confocal (Fig. 5d) imaging can resolve individual fibers and have a significantly reduced background. Figure 5e quantifies the intensities of the plot profiles in the insets of Fig. 5b-d. In the axon region (Fig. 5ei) the confocal and SLM-pick profiles both show three peaks, but the widefield profile only shows two peaks. Likewise, in the dendrite region (Fig. 5eii) the confocal profile (blue) shows four unambiguous peaks corresponding to dendrites near positions 3.3, 5.0, 5.8, and 6.9 μm. The SLM-pick profile (purple) shows the same four peaks with similar relative intensities, but the widefield profile (red) only shows one unambiguous peak. The widefield image also shows a significantly higher background than SLM-pick or confocal images. Near the center dendrite, the background intensity is more than half of the dendrite intensity. In summary, the imaging of tightly-bundled neuronal fibers shows that SLM-pick imaging rejects stray light, improving the contrast of fibers with their backgrounds and allowing them to be clearly discerned, even at submicron resolution.

**Figure 5.**
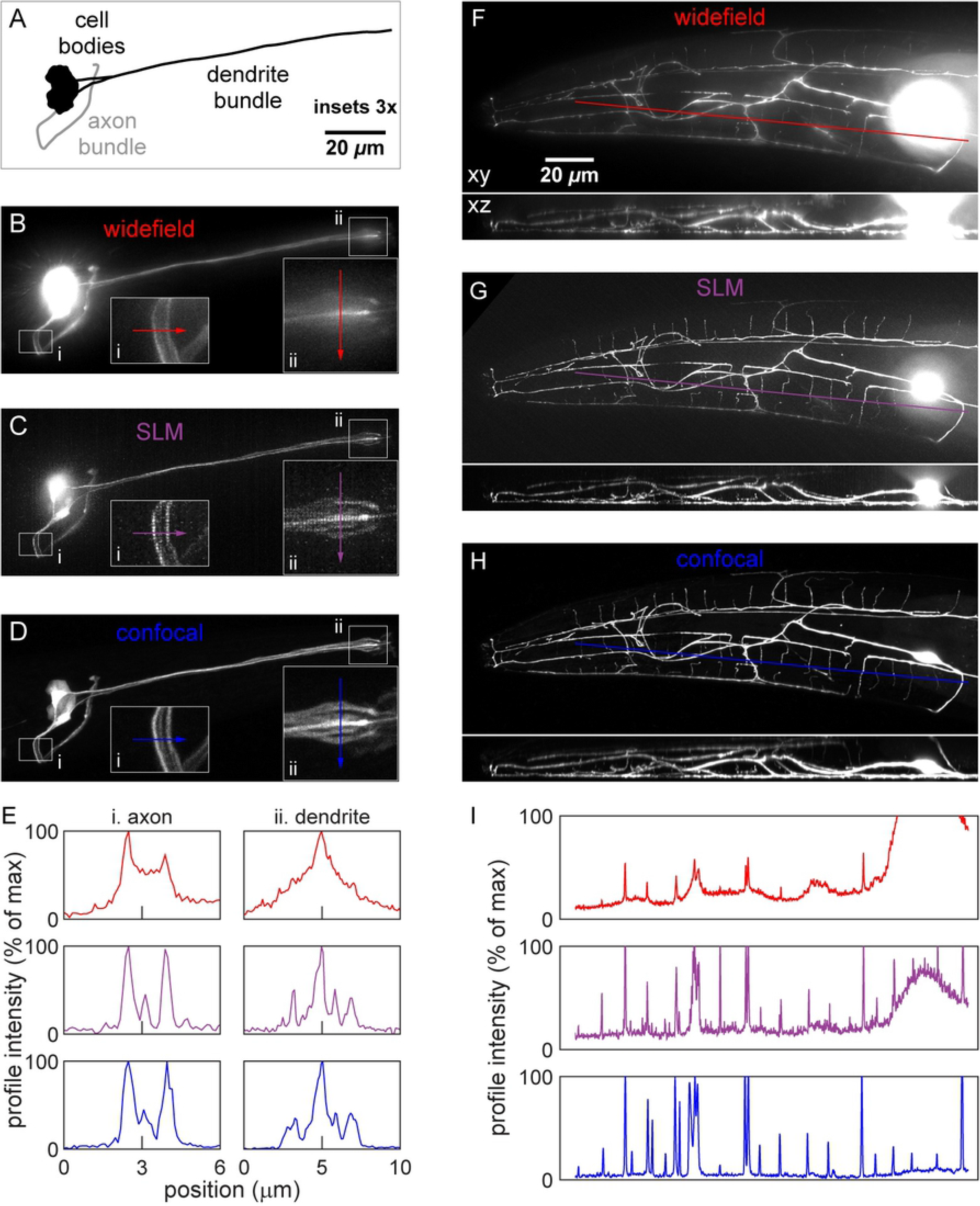
*In vivo* imaging of fluorescent neurons in *C. elegans*. (a) Line drawing of amphid neurons imaged. General location of cell bodies and neuronal fibers shown for simplicity. (b-d) Fluorescence amphid neuron images taken by widefield (b), SLM-pick (c), and confocal (d) microscopes. Insets show 3x expanded view of axons (i) and dendrites (ii) with line profiles. (e) Intensity profile of axons (i) and dendrites (ii). Peaks correspond to clear fibers in fluorescence images (b-d). Note correlation of peaks in SLM and confocal profiles and large background obscuring fibers in widefield profile. (f-h) Fluorescence images of FLP neuron taken by widefield (f), SLM-pick (g), and confocal (h) microscopes. (i) Intensity profile in images (f-h). Note correlation of peaks in SLM and confocal profiles and large background in widefield profile, particularly near cell body.

Second, to demonstrate enhanced imaging capabilities in 3D and at depth, we imaged a *C. elegans* FLP neuron, which has a highly branched 3D dendritic structure [13]. Throughout the extent of the FLP neuron, the SLM-pick images (Fig. 5g) exhibit a better signal-to-background compared to widefield (Fig. 5f), particularly near the cell body, whose contribution to the widefield background is greater than the signal of nearby fibers. The confocal image (Fig. 5h) and profile (Fig. 5i, blue) shows 25 fibers and peaks, respectively, along the path shown. The SLM-pick profile shows 23 peaks, and the widefield profile only shows 11 peaks. The strong correlation of SLM-pick with confocal imaging demonstrates the capability of SLMs to enhance imaging throughout the 3D bulk of the sample. Also, the SLM and confocal imaging show enhanced optical sectioning ability at all depths compared to widefield imaging (see 3D rotation in Supp. Movie S1). This is evidenced by thinner, better-defined dendrites in the xz axial view, especially at the deeper layers (upper side of xz views).

## Discussion

Our study uniquely combines components and techniques of many prior studies and setups. First, there is an effort to spatially modulate the illumination, as reviewed in [2]. In laser-scanning confocal and two-photon microscopy, the laser beam intensity is controlled by electro- or acousto-optic modulators. In widefield imaging, SLMs or DMDs pattern the illumination and can scan samples without macroscopically-moving parts. Depending on the implementation, modulation of the beam can reduce photobleaching and phototoxicity [14], optically section the sample [6], or increase dynamic range [15]. Second, there are efforts to replace the pinhole and PMT of confocal setups by an array camera [16–18]. One technique that is very similar to ours is called Image Scanning Microscopy, which rescans the emission beam of a confocal microscope after the pinhole and utilizes an camera array [16, 19]. Several studies use more complicated postprocessing, such as Gaussian masks and deconvolution to achieve confocal capability in widefield imaging [11, 20]. The studies above lay the foundation for our study, which details one of the simplest and most cost-effective methods for converting a widefield into a confocal microscope. The cost of confocal microscopes often renders them inaccessible to individual laboratories. In our setup, a single SLM ($70) mounted onto a high-precision stage ($500) are the only add-ons required to give confocal capabilities to a ubiquitous instrument.

In this study, we utilized an SLM to modulate the light distribution at the field stop of our inverted microscope to project an arbitrary illumination pattern on the focal plane in the sample. We show that the SLM-max postprocessing produces images with optical sectioning similar to SLM-pick postprocessing while relaxing requirements for alignment and stability in space and time. Utilizing this setup with SLM-pick postprocessing, we rigorously characterized optical sectioning of well-defined samples, including fluorescent thin film and beads. Compared to widefield imaging, we demonstrate enhanced optical sectioning and improved signal-to-background at ratio high resolution and at depth in fluorescent neurons *in vivo*.

For demonstration purposes, we utilized an inexpensive, off-the-shelf SLM with limited characteristics (*e.g.*, transmission, fill factor, and element size). As a result, we experienced challenges in obtaining adequate emission light for imaging deep in some samples and observed pixelation in the resulting images. Out-of-focus fluorescence from bright objects, such as the cell body, also remains in our SLM-pick images because our off-the-shelf SLM has larger elements, necessitating a larger virtual pinhole and reduced optical sectioning. Even so, our images suffice to demonstrate optical sectioning and enhanced imaging. With an improved SLM or by utilizing a DMD, which has superior specifications, we expect improved optical sectioning and resolution with better contrast and deeper imaging.

## Materials and methods

### SLM setup

Following a prior study [8], we removed an SLM from a digital projector (Epson PowerLite 1810p). The SLM was placed between crossed polarizers (Edmund Linear Polarizing Film XP42-18), aligned by hand using a power meter. We measured a 200:1 extinction ratio (transmission of element when on to off). We used a 6-axis stage (Newport, model 9031) and a custom 3D-printed mount to hold and align the SLM. We used Matlab 2020a (MathWorks) to control transmission through the SLM elements. The code is available at Github (https://github.com/wormneurolab/SLM-confocal).

### SLM alignment

Utilizing the center row and column elements of the SLM, we projected an image of a cross onto a fluorescent thin film. We adjusted the SLM position and orientation using the 6-axis stage so that the cross was in focus and centered on the camera array. This alignment took about a few minutes and was stable for a day.

### SLM illumination and imaging

As detailed in Fig. 3ab, utilizing the MATLAB to control the SLM we illuminated the sample with 36 raster-scanned dot array patterns and captured 36 raw sub-images sequentially. Each sub-image contains information from the full camera array with 1/36^th^ of the FOV illuminated. The exposure time depends on intensity of the light source, transmission of the SLM, strength of fluorescent labelling, and sensitivity of the camera. Utilizing our setup and samples, we acquired all the raw data for a 2D image in about 5 seconds. This extended time was primarily due to the low transmission of our SLM. We expect significantly reduced exposure time with an improved SLM or a DMD.

### Postprocessing

The SLM-sum and SLM-max strategies produce a final image where the pixel values are the sum and the maximum values, respectively, of the corresponding pixels in the 36 sub-images. The SLM-max strategy is commonly known as a maximum projection. In the SLM-pick strategy the values of each pixel in the final image are the values in the sub-image when the pixels are illuminated. Because the pixel-to-element ratio is not an integer (~3.5) some pixels are partially illuminated by two SLM elements. For those pixels, we used the maximum value of the pixels in the sub-images.

### Fluorescent thin film

Using a fluorescent highlighter pen, we drew a 3-mm diameter spot on a slide. We covered the spot with a coverslip and applied pressure, generating a thin film between coverslip and slide. We taped two edges of the thin film slide and allowed it to dry at room temperature for 4 hours.

### Fluorescent bead

For 6-μm beads, we centrifuged 50 μL of 10% bead solution (Invitrogen I14785) for 1 min, removed 25 μL of the supernatant, and vortexed the remaining 25 μL solution for 5 mins. We dropped 5 μL of the concentrated solution to the coverslip center, then added 5 μL to the same location to increase bead density. We dried the coverslip at 37 °C for 5 mins. Then we dropped 7 μL of mounting solution to a slide center, flipped over the coverslip, and placed it on the slide without shearing movement. We dried the slide at 37 °C for 15 mins and sealed the coverslip edges with wax. For 175-nm beads (Molecular Probes P7220), we followed established procedures [21].

### Microscopy

We used widefield, SLM-enabled, and confocal microscopy to take 3D image stacks (*i.e.*, z-stack). We utilized a Nikon Ti2-E inverted microscope with a SOLA SE II LED light engine and a 1.4 NA, 60x objective for widefield and SLM-enabled imaging. We utilized a Zeiss LSM 800 microscope with a 1.4 NA, 63x objective for confocal imaging. For thin films, we imaged a 6-μm depth with 50-nm step size. For 6-μm beads, we imaged a 20-μm depth with 250-nm step size. For 175-nm beads, we imaged a 6-μm depth with 50-nm step size. We describe animal imaging below.

### Image analysis

We averaged thin film and bead data by aligning multiple measurements. For thin film, we used the z position of maximum brightness (found by Gaussian fitting) to align and average profiles together. For 6-μm fluorescent beads, we located each bead and found z position of its center using the ImageJ [22] plugin “3D Objects Counter” (https://imagej.net/3D_Objects_Counter). Using the nearest z slice image, we employed a 2D Gaussian to fit the bead image in x and y. Thus, we obtained the bead’s center position, and utilizing this position, we averaged bead images together and generated transverse and axial profiles. For 175-nm beads, we followed established procedures [21] utilizing the analysis software PSFj [23] to measure the point spread function to assess resolution.

*C. elegans* cultivation, immobilization, imaging: We followed established procedure for *C. elegans* strains cultivation on Bacto agar plates [24] at 15°C, animal immobilization by sodium azide, and imaging [25]. After immobilization, animals were rotated to a desired orientation [26] under a fluorescence stereomicroscope and then imaged under an inverted microscope.

---

**Figure S1. Rectangular active region of SLM.** Brightfield image of SLM between two crossed polarizers. Bright areas are rectangular active regions of SLM elements. Measurements given in pixels of brightfield microscope camera.

**Figure S2. SLM-sum postprocessing strategy.** Fluorescence images obtained by maximum projections of 3D image. (a) Conventional widefield image, replicated from Fig. 3d. (b) SLM-sum image. Pixel value in image is sum of pixel values in 36 sub-images.

**Table S1. Transverse and axial resolution of widefield, SLM-max, and SLM-pick imaging.** FWHM of 175-nm fluorescent beads. SLM-max and SLM-pick do not show improvement of transverse or axial resolution compared to widefield, due to illumination area larger than diffraction limit.

**Movie S1. FLP neuron imaged by widefield, SLM-pick, and confocal microscopy.** Movie shows 3D rendering of neuron in Fig. 5f-h.

## Acknowledgements

Some nematode strains used in this work were provided by the Caenorhabditis Genetics Center (CGC), which is funded by the NIH National Center for Research Resources (NCRR). NG3146 and CHB1226 are gifts from Gian Garriga and Maxwell Heiman, respectively. We acknowledge Kai Zhang from Bryan Spring’s laboratory (Northeastern Physics Dept.) for helpful discussion in resolving SLM aliasing. We acknowledge Noah Joseph (Northeastern Bioengineering Dept.) for 3D printing the SLM mount. The authors were supported in part by a Northeastern University TIER 1 grant.

## References

1. Pawley J. Handbook of Biological Confocal Microscopy: Springer US; 2013.

2. Krishnaswami V, Van Noorden CJF, Manders EMM, Hoebe RA. Spatially-controlled illumination microscopy. Q Rev Biophys. 2016;49:e19. Epub 2016/12/12. doi: 10.1017/S0033583516000135.

3. Nikolenko V, Peterka DS, Yuste R. A portable laser photostimulation and imaging microscope. J Neural Eng. 2010;7(4):7. doi: 10.1088/1741-2560/7/4/045001. PubMed PMID: WOS:000280038600002.

4. Yang W, Jae E, Carrillo-Reid L, Pnevmatikakis E, Paninski L, Yuste R, et al. Simultaneous Multi-plane Imaging of Neural Circuits. Neuron. 2016;89(2):269–84. doi: 10.1016/j.neuron.2015.12.012.

5. Rector DM, Ranken DM, George JS. High-performance confocal system for microscopic or endoscopic applications. Methods. 2003;30(1):16–27. doi: 10.1016/s1046-2023(03)00004-5. PubMed PMID: WOS:000182506200003.

6. Hanley QS, Verveer PJ, Gemkow MJ, Arndt-Jovin D, Jovin TM. An optical sectioning programmable array microscope implemented with a digital micromirror device. J Microsc-Oxf. 1999;196:317–31. PubMed PMID: WOS:000084436500006.

7. Fiolka R, Beck M, Stemmer A. Structured illumination in total internal reflection fluorescence microscopy using a spatial light modulator. Opt Lett. 2008;33(14):1629–31. doi: 10.1364/ol.33.001629. PubMed PMID: WOS:000258309900027.

8. Huang D, Timmers H, Roberts A, Shivaram N, Sandhu AS. A low-cost spatial light modulator for use in undergraduate and graduate optics labs. Am J Phys. 2012;80(3):211––5. doi: 10.1119/1.3666834. PubMed PMID: 820e79f8ae7b4dbe96c8005e80761bb2.

9. Davidovits P, Egger MD. Scanning Laser Microscope. Nature. 1969;223(5208):831-. doi: 10.1038/223831a0.

10. Petráň M, Hadravský M, Egger MD, Galambos R. Tandem-Scanning Reflected-Light Microscope. J Opt Soc Am. 1968;58(5):661–4. doi: 10.1364/JOSA.58.000661.

11. York AG, Parekh SH, Nogare DD, Fischer RS, Temprine K, Mione M, et al. Resolution doubling in live, multicellular organisms via multifocal structured illumination microscopy. Nat Methods. 2012;9(7):749–U167. doi: 10.1038/nmeth.2025. PubMed PMID: WOS:000305942200034.

12. Wilhelm S, Grobler B, Gluch M, Heinz H. Confocal Laser Scanning Microscopy. Principles. Microscopy from Carl Zeiss. 2003.

13. Smith CJ, Watson JD, Spencer WC, O’Brien T, Cha B, Albeg A, et al. Time-lapse imaging and cell-specific expression profiling reveal dynamic branching and molecular determinants of a multi-dendritic nociceptor in C. elegans. Dev Biol. 2010;345(1):18–33. Epub 2010/06/09. doi: 10.1016/j.ydbio.2010.05.502. PubMed PMID: 20537990.

14. Hoebe RA, Van Oven CH, Gadella TWJ, Dhonukshe PB, Van Noorden CJF, Manders EMM. Controlled light-exposure microscopy reduces photobleaching and phototoxicity in fluorescence live-cell imaging. Nat Biotechnol. 2007;25(2):249–53. doi: 10.1038/nbt1278. PubMed PMID: WOS:000244064000032.

15. Yang RH, Weber TD, Witkowski ED, Davison IG, Mertz J. Neuronal imaging with ultrahigh dynamic range multiphoton microscopy. Sci Rep. 2017;7:7. doi: 10.1038/s41598-017-06065-7. PubMed PMID: WOS:000405894200035.

16. Sheppard CJR. Super-resolution in Confocal Imaging. Optik. 1988;80(2):53–4. PubMed PMID: WOS:A1988Q132800002.

17. Roth S, Sheppard CJR, Wicker K, Heintzmann R. Optical photon reassignment microscopy (OPRA). Optical Nanoscopy. 2013;2(1):5. doi: 10.1186/2192-2853-2-5.

18. De Luca GMR, Breedijk RMP, Brandt RAJ, Zeelenberg CHC, de Jong BE, Timmermans W, et al. Re-scan confocal microscopy: scanning twice for better resolution. Biomed Opt Express. 2013;4(11):2644–56. doi: 10.1364/boe.4.002644. PubMed PMID: WOS:000326582500031.

19. Muller CB, Enderlein J. Image Scanning Microscopy. Phys Rev Lett. 2010;104(19):4. doi: 10.1103/PhysRevLett.104.198101. PubMed PMID: WOS:000277699600054.

20. Heintzmann R, Benedetti PA. High-resolution image reconstruction in fluorescence microscopy with patterned excitation. Appl Opt. 2006;45(20):5037–45. doi: 10.1364/ao.45.005037. PubMed PMID: WOS:000239018000032.

21. Cai H, Wang YL, Wainner RT, Iftimia NV, Gabel CV, Chung SH. Wedge prism approach for simultaneous multichannel microscopy. Sci Rep. 2019;9(1):17795. doi: 10.1038/s41598-019-53581-9. PubMed Central PMCID: PMCPMC6882912.

22. Schindelin J, Arganda-Carreras I, Frise E, Kaynig V, Longair M, Pietzsch T, et al. Fiji: an open-source platform for biological-image analysis. Nat Methods. 2012;9(7):676–82. doi: 10.1038/nmeth.2019. PubMed PMID: WOS:000305942200021.

23. Theer P, Mongis C, Knop M. PSFj: know your fluorescence microscope. Nat Meth. 2014;11(10):981–2. doi: 10.1038/nmeth.3102.

24. Brenner S. The genetics of Caenorhabditis elegans. Genetics. 1974;77(1):71–94. PubMed PMID: 4366476.

25. Chung SH, Clark DA, Gabel CV, Mazur E, Samuel AD. The role of the AFD neuron in C. elegans thermotaxis analyzed using femtosecond laser ablation. BMC Neurosci. 2006;7:30. PubMed PMID: 16600041; PubMed Central PMCID: PMCPMC1450292.

26. Chung SH, Mazur E. Femtosecond laser ablation of neurons in C. elegans for behavioral studies. Appl Phys A-Mater Sci Process. 2009;96(2):335–41. PubMed PMID: ISI:000267095400007; PubMed Central PMCID: PMCN/A.

